# Characterising the short-term habituation of event-related evoked potentials

**DOI:** 10.1101/153387

**Authors:** Flavia Mancini, Alessia Pepe, Alberto Bernacchia, Giulia Di Stefano, André Mouraux, Gian Domenico Iannetti

**Author notes:** Corresponding author:* Flavia Mancini University of Cambridge Department of Engineering, Computational and Biological Learning Lab Trumpington Street, Cambridge, CB2 lPZ.

## Abstract

Fast-rising sensory events evoke a series of functionally heterogeneous event-related potentials (ERPs). Stimulus repetition at 1 Hz is known to induce a strong habituation of the largest ERP responses, the vertex waves, which are elicited by stimuli regardless of their modality^7^, provided that they are salient and behaviourally-relevant. In contrast, the effect of stimulus repetition on the earlier sensor)^7^ components of ERl’s has been less explored, and the few existing results are inconsistent. To characterize how the different ERP waves habituate over time, we recorded the responses elicited by 60 identical somatosensory stimuli (activating either non-nociceptive **A** β or nociceptive A5 afferents), delivered at 1 Hz to healthy human participants. We show that the well-described spatiotemporal sequence of lateralised and vertex ERP components elicited by the first stimulus of the series is largely preserved in the smaller-amplitude, habituated response elicited by the last stimuli of the series. We also found that the earlier lateralised sensory waves habituate across the 60 trials following the same decay function of the vertex waves: this decay function is characterised by a large drop at the first stimulus repetition followed by smaller decreases at subsequent repetitions. Interestingly, the same decay functions described the habituation of ERPs elicited by repeated non-nociceptive and nociceptive stimuli. This study provides a neurophysiological characterization of the effect of prolonged and repeated stimulation on the main components of somatosensory ERPs. It also demonstrates that both lateralised waves and vertex waves are obligator}^7^ components of ERPs elicited by non-nociceptive and nociceptive stimuli.

**Significance statement:** Our results provide a functional characterization of the decay of the different ERP components when identical somatosensory (nociceptive and non-nociceptive) stimuli are repeated at 1Hz. Fast-rising stimuli elicit ERPs obligator)^7^ contributed by both early lateralised components and late vertex components, even when stimulus repetition minimizes stimulus relevance. This challenges the view that lateralised waves are not obligatorily elicited by nociceptive stimuli. Furthermore, the lateralised and vertex waves habituate to stimulus repetition following similar decay functions, which are unlikely explained in terms of fatigue or adaptation of skin receptors.

## Introduction

Sudden sensory events evoke a series of transient responses in the ongoing electrocortical activity (event-related potentials, ERPs). ERPs are functionally heterogeneous and reflect the activity of distinct cortical generators overlapping in time and space (Sutton et al., 1965). Since these generators include both sensory and associative cortical areas, the scalp distribution of the early lateralised ERP components elicited by stimuli of different modalities partly differs depending on the modality of the sensor)^7^ input. In contrast, the scalp distribution of the late and largest ERP components is virtually identical regardless of the modality of the eliciting stimulus (Mouraux and Iannetti, 2009): it consists in a biphasic negative-positive deflection widespread over the scalp and maximal at the vertex — often referred to as Vertex wave’ or vertex potential’ (Bancaud et al., 1953).

The vertex wave amplitude is maximal when fast-rising stimuli are presented using large and variable inter-stimulus intervals of several seconds (Mouraux and Iannetti, 2009; Huang et al., 2013), or when the stimulus reflects behaviourally relevant changes within a regular series of otherwise identical stimuli (Snyder and Hillyard, 1976; Valentini et al., 2011; Ronga et al., 2013). In contrast, when identical stimuli are monotonously repeated at short and regular intervals (e.g., 0.5 or 1 Hz), the vertex wave amplitude strongly decays (jasper and Sharpless, 1956; Ritter et al., 1968; Davis et al., 1972; Mouraux and lannetti, 2009; Liang et al., 2010; Wang et al., 2010). Although the decay of the vertex wave due to repeated stimulation at different frequencies has been described (Fruhstorfer et al., 1970; Greffrath et al., 2007), a formal characterization of how the different constituent components of the ERP habituate over time is still missing. This is particularly important considering that previous studies suggested that neural activity in different cortical regions adapts to repeated stimulation at different timescales: for instance, neural activity in associative regions elicited by trains of innocuous, somatosensory stimuli decays faster than neural activity in sensory cortices (Forss et al., 2001; Venkatesan et al., 2014). However, these results may not generalise to responses elicited by noxious somatosensory stimuli: a previous study has suggested that the repetition of intra-epidermal nociceptive stimuli at 1 Hz for 1 minute fully suppresses lateralized evoked responses (Mouraux et al., 2013).

Therefore, our primary objective was to describe the short-term habituation of the different constituents of somatosensory nociceptive and non-nociceptive ERl’s: both the large centrally-distributed vertex waves (N2 and 1’2 waves) and the smaller lateralised somatosensory waves (N1 and 1’4 waves). These are all the known waves elicited by nociceptive stimulation (Treede et al., 1988; Valentini et al., 2012; Hu et al., 2014). As in Mouraux et al. (2013), we recorded EEG while delivering trains of 60 identical stimuli at 1 Hz. In one group of healthy participants, we transcutaneously and electrically stimulated nerve trunks, activating directly all large-diameter A β somatosensory afferents and eliciting non-painful sensations. In a separate group of participants, we used radiant-heat stimuli that selectively activate skin nociceptors and elicit sensations of Aδ-mediated pinprick pain. We did not use intra-epidermal electrical stimulation of nociceptive afferents (Mouraux et al., 2013), because it can induce strong habituation of peripheral nociceptors (the stimulus is delivered always in the same location, whereas radiant heat stimuli can be easily displaced to reduce nociceptor fatigue). The use of two different somatosensory stimuli allowed to cross-validate and generalise our findings across different sensor) pathways.

We addressed two complementary questions. (1) First, we statistically assessed whether the main response components were present in both the non-habituated ERP (i.e. the ERP elicited by the first stimulus of a series) and the habituated ERP (i.e. the ERP elicited by later stimuli that elicit a stable, habituated response). The rationale for this decision was the consistent observation that the amplitude of the main ERP waves (i.e., vertex waves) decays only minimally after the first few stimulus repetitions (Ritter et al., 1968; Fruhstorfer et al., 1969; Fruhstorfer et al., 1970; Fruhstorfer, 1971; Greffrath et al., 2007; Mouraux et al., 2013), a finding corroborated by the present results (Figure. 1-4). (2) Second, we asked whether and how the lateralized and vertex waves habituated throughout the block of 60 stimuli. We used Singular Value Decomposition (SVD) to separate the ERP waveform from its amplitude change across stimulus repetitions. SVD provides a small number of components that best approximate the data and explain most of its variance (Golub and Reinsch, 1970). This approach allowed us to investigate the decay function of small ERP components, such as the lateralized waves.

**Figure 1.**
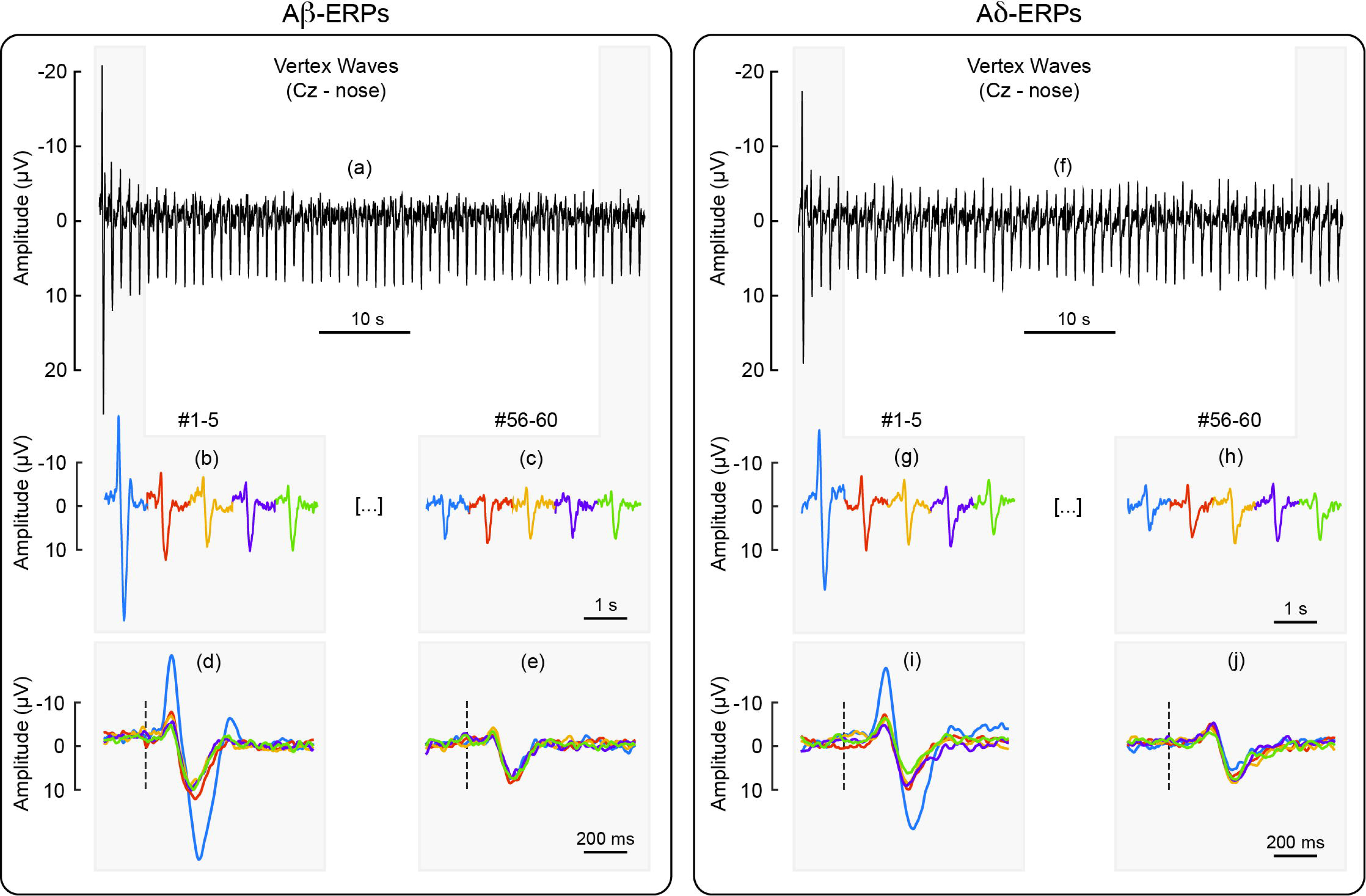
Habituation of vertex waves (N2, P2) elicited by repeated Aβ (panels a-e) and Aδ stimuli (panels f-j), at electrode CZ referenced to the nose. Panel a shows the vertex waves elicited by 60 A β stimuli delivered at 1 Hz, whereas panel f shows the vertex waves elicited by 60 Aδ stimuli delivered at the same frequency. To facilitate visual comparison, the figure displays, as enlarged and concatenated, the responses to the first five Aβ stimuli (panel b), the last five Aβ stimuli (panel c), the first five Aδ stimuli (panel g), and the last five Aδ stimuli (panel h). The figure also displays, as enlarged and super-imposed, the same responses to the first five A β stimuli (panel d), the last five Aβ stimuli (panel e), the first five Aδ stimuli (panel i), and the last five Aδ stimuli (panel j).

## Methods

### Participants

Thirty-two healthy subjects (14 women) aged 19—31 years (mean ± SD: 23.6 ± 3.9) participated in the study, after having given written informed consent. All experimental procedures were approved by the ethics committee of University^7^ College London (2492/001).

### Transcutaneous electrical stimulation of Aβ fibers

Innocuous stimulation of Aβ afferents consisted of square-wave pulses (100 µs duration), generated by a constant current stimulator (DS7A, Digitimer, UK). Stimuli were delivered through a bipolar electrode placed above the superficial radial nerve and elicited a paresthetic sensation in the corresponding innervation territory. Aβ detection thresholds were identified using the method of ascending staircases, on the right hand. The detection threshold was defined as the average of the lowest stimulus energy eliciting a sensation in 3 consecutive trials. Electrical stimuli were delivered at approximately 300% of each individual’s Aβ detection threshold. Stimulus intensity was slightly adjusted to elicit sensations of comparable intensities on the left and right hands (mean ± SD, 17.4 ± 11.4 mA) and to make sure that the elicited sensation was never painful.

### Cutaneous laser stimulation of Aδ and C fibers

Nociceptive stimuli were radiant heat pulses generated by an infrared neodymium:yttrium-aluminum-perovskite laser with a wavelength of 1.34 µm (Nd:YAP; Electronical Engineering, Italy). At this wavelength, laser pulses excite Aδ and C nociceptive free nerve endings in the epidermis directly and selectively, i.e. without coactivating touch-related Aβ fibers in the dermis (Bromm and Treede, 1984; Baumgartner et al., 2005; Mancini et al., 2014). The duration of each laser pulse was 4 ms.

Laser stimuli were delivered within a squared skin area (4×4 cm) centered on the dorsum of the hand, encompassing the area in which the stimulation of Aβ afferents elicited the paraesthesia. The laser beam was transmitted through an optic fiber, and its diameter at target site was set at ~6 mm by focusing lenses. A visible He—Ne laser pointed to the stimulated area, within which the laser beam was manually displaced after each stimulus. The laser was triggered by a computer script.

The method of ascending staircases used for identifying the detection threshold of Aβ stimuli was also used to identify the detection threshold of Aδ stimuli. For the EEG recordings, the stimulus energy was clearly above the activation threshold of A5 fibers (0.53 + 0.06 J/mm^2^). This stimulus energy elicited intense but tolerable pinprick pain sensations, of comparable intensities on the right and left hands. Because variations in baseline skin temperature may modulate the intensity of the afferent nociceptive input (Iannetti et al., 2004), an infrared thermometer was used to ensure that the hand temperature varied no more than 1°C across blocks. To avoid receptor fatigue or sensitization, the laser beam was displaced after each stimulus by ~1 cm within the predefined stimulated area.

### Experimental procedure

Participants sat comfortably with their hands resting on a table in front of them. They were instructed to focus their attention on the stimuli and fixate a yellow circular target (diameter: 1 cm) placed in front of them at a distance of approximately 60 cm from their face. A black curtain blocked the view of the hands. Throughout the experiment, white noise was played through headphones, to mask any sound associated with the either type of somatosensory stimulation.

The experiment was performed on 32 participants, divided in two groups of 16 participants. One group received electrical stimuli, and the other group received laser stimuli, using an identical procedure. Each participant received the somatosensory stimuli in 10 blocks, separated by a 5-minute interval, during which participants were allowed to rest. Each block consisted of 60 somatosensory stimuli delivered at 1 Hz: thus, each block lasted 1 minute. In each block, stimuli were delivered either to the right hand or to the left hand. Right- and left-hand blocks were alternated. The order of blocks was balanced across participants; half of the subjects started with a right-hand block, and the other half started with a left-hand block. At the end of each block, participants were asked to provide an average rating of perceived stimulus intensity, with reference to the modality of the stimulus and using a numerical scale ranging from 0 (“no shock sensation” or “no pinprick sensation”) to 10 (“most intense shock sensation” or “most intense pinprick sensation”). This was done to ensure that the perceived intensity of the stimuli was similar across blocks (rating variability, SD across blocks: electrical stimuli, 0.2 ± 0.2; laser stimuli: 3 ± 0.4).

### Electrophysiological recordings

EEG was recorded using 30 Ag—AgCl electrodes placed on the scalp according to the International 10-20 system (Electro-Cap International; USA), using the nose as reference. Electrode positions were T’pT, ‘Fpz’, ‘Fp2’, ‘F7’, ‘F3’, ‘Fz’, ‘F4’, ‘F8’, T3’, ‘C3’, ‘Cz’, ‘C4’, T4’, T5’, T3’, ‘Pz’, T4’, T6’, ‘Ol’, ‘Oz’, ‘02’, ‘FCz’, ‘FC4’, ‘FC3’, ‘Cp3’, ‘Cp4’. Eye movements and blinks were recorded from the right *orbicularis oculi* muscle, using 2 surface electrodes. The active electrode was placed below the lower eyelid, and the reference electrode a few centimetres laterally to the outer canthus. Signals were amplified and digitized using a sampling rate of 1,024 Hz (SD32; Micromed, Italy).

### EEG analysis

*1. Preprocessing.* EEG data were preprocessed and analyzed using Letswave 6 and EEGLAB (https://sccn.ucsd.edu/eeglab/). Continuous EEG data were band-pass filtered from 0.5 to 30 Hz using a Butterworth filter, segmented into epochs using a time window ranging from −0.2 to 0.8 sec relative to the onset of each stimulus, and baseline corrected using the interval from −0.2 to 0 sec as reference. Trials contaminated by large artefacts (<10% per condition) were removed. Eye blinks and movements were corrected using a validated method based on unconstrained Independent Component Analysis (“runica” algorithm of EEGLAB). In all datasets, independent components related to eye movements showed a large EOG channel contribution and a frontal scalp distribution. To allow averaging across blocks while preserving the possibility of detecting lateralized EEG activity, scalp electrodes were flipped along the medio-lateral axis for all signals recorded in response to left hand stimulation. Hereinafter, we refer to the central electrode contralateral to the stimulated hand as Cc. In each participant, we averaged each of the 60 ERl’ responses across the 10 recording blocks, and thus obtained 60 average ERl’ waveforms: one for each of the 60 trials and for each participant.

*2. Statistical assessment of ERP components.* We assessed the consistency of stimulus-evoked modulations of EEG amplitude across time, to statistically evaluate whether EEG deflections in the post-stimulus time window (from 0 to +0.8 s) was significantly greater than baseline. Specifically, we performed a one-sample t-test against zero (i.e. against baseline) for each electrode and time point of the entire baseline-corrected, single-subject waveforms, using cluster-level permutation testing. This analysis yielded a scalp distribution of t-values across time and was performed separately on the non-habituated ERl’ and on the habituated ERP of each modality.

The non-habituated ERl’ was derived, for each participant, by averaging all the responses elicited by the 1^st^ stimulus of all blocks. The habituated ERP was derived, for each participant, by averaging the responses elicited by the 6^th^ to the 60^th^ stimuli of all blocks. The decision of using these responses elicited by stimuli 6^th^ to 60^th^ as a proxy of the habituated ERP was based on the observation that the amplitude of the main ERP waves decays only minimally after the first 5 stimulus repetitions, as observed here (Figure 1-2, 4) and previously described (Fruhstorfer et al., 1970; Greffrath et al., 2007). Figure. 1 and 2 show how the amplitude of the ERPs was consistently habituated after the first few stimulus repetitions.

**Figure 2.**
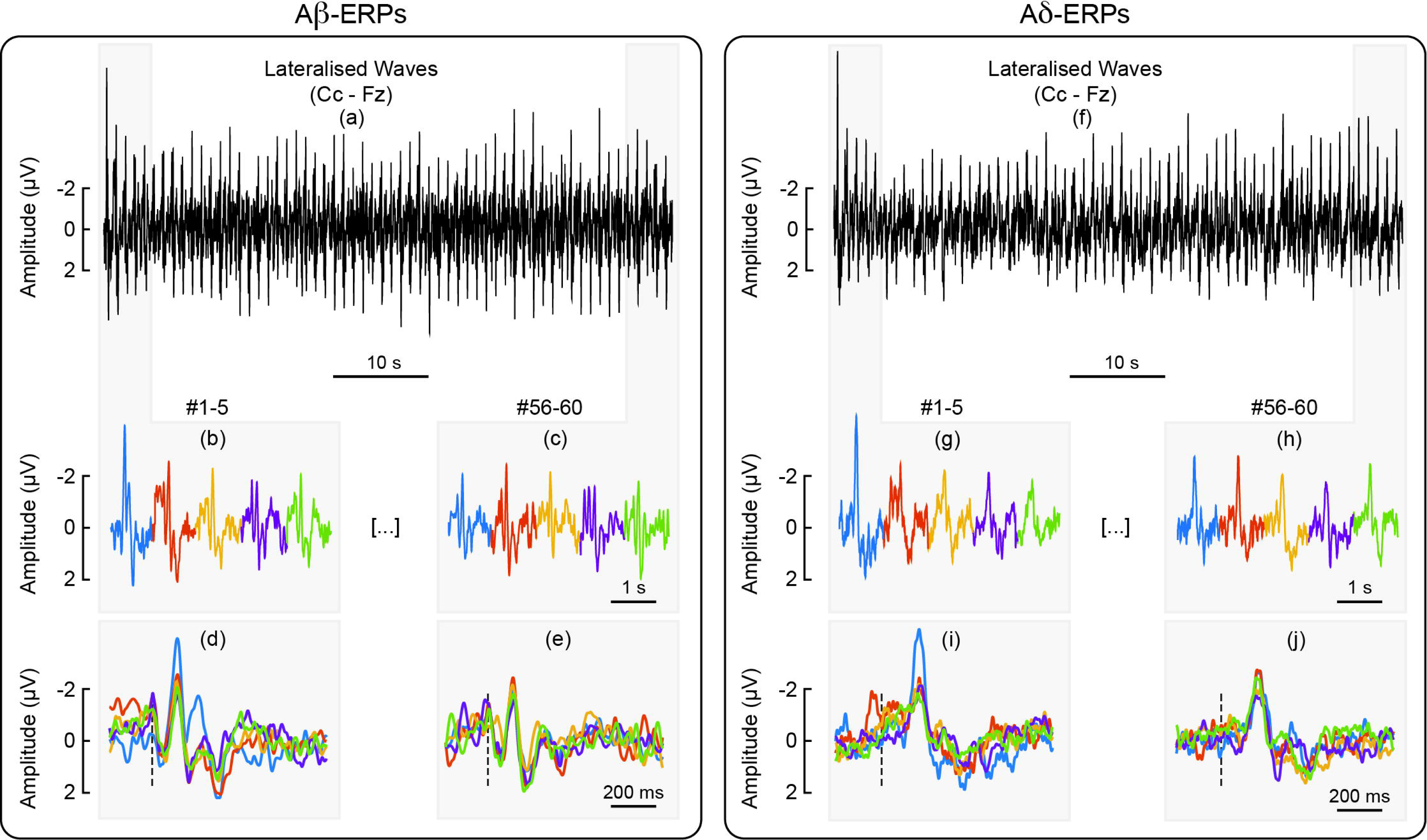
Habituation of lateralised somatosensory waves (N1, P4) elicited by repeated Aβ (panels a-e) and Aδ stimuli (panels f-j), at the central electrode contralateral to hand stimulation (Cc) referenced to the nose. Panel a shows the lateralized waves elicited by 60 A β stimuli delivered at 1 Hz, whereas panel f shows the lateralized waves elicited by 60 Aδ stimuli delivered at the same frequency. To facilitate visual comparison, the figure displays, as enlarged and concatenated, the responses to the first five Aβ stimuli (panel b), the last five Aβ stimuli (panel c), the first five Aδ stimuli (panel g), and the last five Aδ stimuli (panel h). The figure also displays, as enlarged and super-imposed, the same responses to the first five Aβ stimuli (panel d), the last five Aβ stimuli (panel e), the first five Aδ stimuli (panel i), and the last five Aδ stimuli (panel j).

To account for multiple comparisons, significant time points (p < 0.05) were clustered based on their temporal adjacency (cluster-level statistical analysis). For each cluster, we calculated the pseudo-/ statistic of the two conditions, estimated its distribution by permutation testing (1000 times), and generated the bootstrap*p* values for testing the null hypothesis that there were no differences in signal amplitude (Maris and Oostenveld, 2007). This procedure identified the clusters in which the response was significantly different than baseline.

T-tests assume that the examined data are normally distributed. To ascertain this, we extracted single-subject peak amplitude values of the components of interest (N1, N2, P2 waves) in each experimental condition, and tested whether they violated normality assumptions using the Shapiro-Wilk test. We did not do this for the 1’4 wave because its detection can be ambiguous in some subjects (Hu et al., 2014), especially in the habituated response. We found moderate-to-strong evidence for normality violation for the A5-P2 peak amplitude at stimuli #1 (p = 0.005) and #6-60 (p = 0.02). We found no evidence of violation to normality distribution for all other waves (p > 0.05). To address the two instances of normality violations, we performed a non-parametric one-sample test (Wilcoxon signed rank test) on the A5-P2 peak values for conditions “stimulus #1” and “stimuli #6-60”, in addition to the point-by-point t-statistics (Gibbons and Chakraborti, 2011). Both tests provided strong evidence that the Aδ-P2 peak values were greater than baseline (stimulus 1: z = 3.52, p < 0.001; stimuli 6-60: z = 3.52, p < 0.001), confirming the results of the point-by-point t-tests reported in Figure 3.

**Figure 3.**
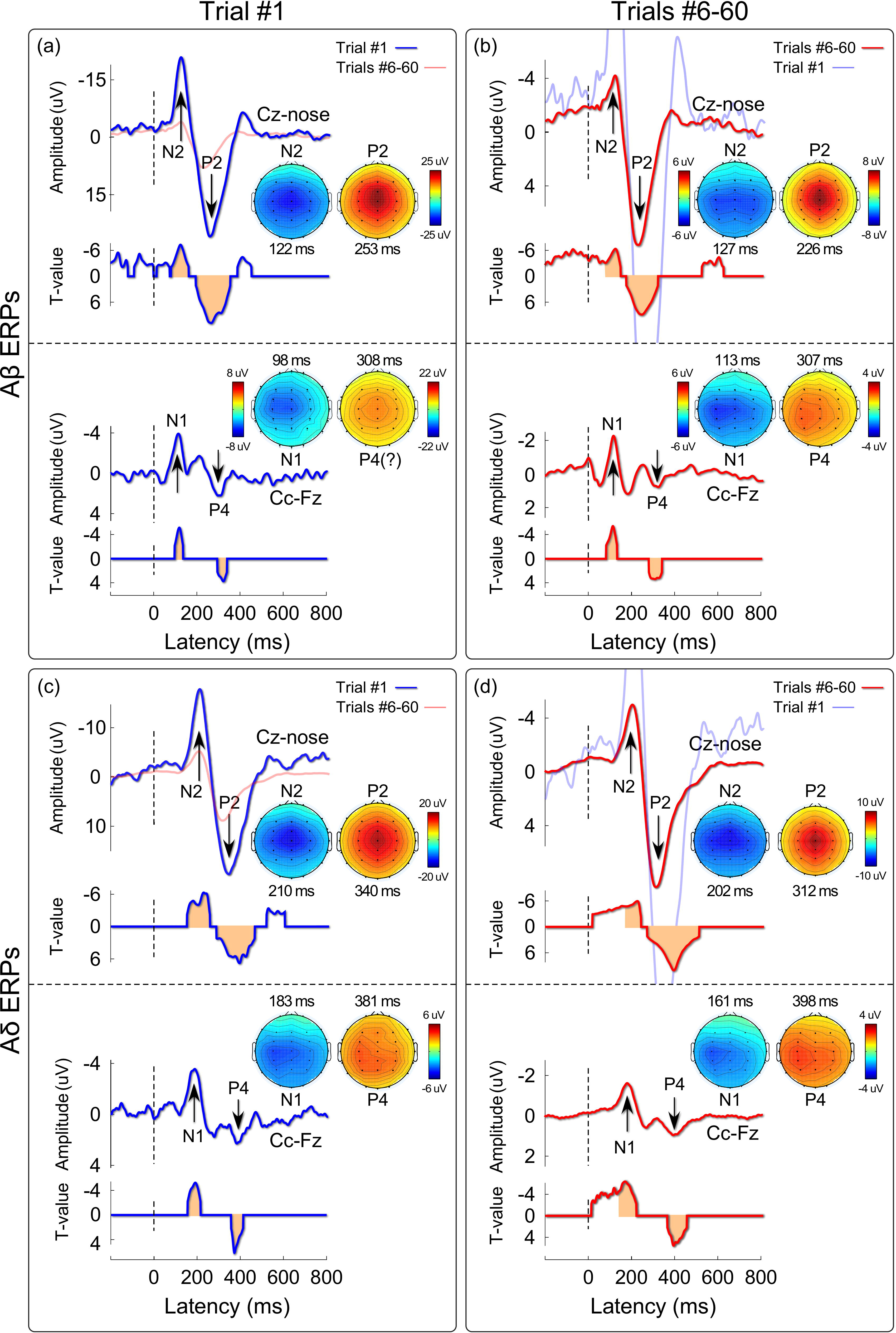
Habituation of vertex waves (N2, P2) and lateralised responses (N1, P4) elicited by Aβ (panels: a, b) and Aδ (panels: c, b) somatosensory stimuli. Displayed signals show group-level ERl’s recorded from the vertex (Cz vs nose) and from the central electrode contralateral to the stimulated hand (Cc vs Ez), elicited by the first stimulus in a series (non-habituated response; panels a, c) and by the average of trials #6-60 (habituated response; panels b, d). Scalp topographies (signals referenced to the nose) are displayed at the peak latency of the Nl, N2, 1’2, and 1’4 waves, in all conditions. The Nl, N2, and 1’2 waves were significantly greater than baseline both in trial #1 and in trials #6-60, as shown by the point-by-point, one-sample *t* statistics plotted below each ERP wave. Time intervals during which the ERP waves were significantly different than 0 in the Nl, N2,1’2, and 1’4 time windows are highlighted in orange.

*3. Modelling the within-block decay of the lateralised and vertex waves.* We tested whether the amplitude of the vertex waves and of the lateralized wave evoked by Aβ and Aδ stimuli was modulated as a function of stimulus repetition. In each participant, we first averaged each of the 60 ERP responses across the 10 recording blocks, and thus obtained 60 average ERP waveforms: one for each of the 60 trials. Then, we averaged across participants and, for each modality, we obtained 60 group-level averages. To study the amplitude modulation of the entire waveform across 60 trials, we decomposed the EEG signals at electrodes of interest (Cz and Cc) using singular-value decomposition (SVD) (Golub and Reinsch, 1970). We used SVD to decompose the modulation of the EEG amplitude across the 1000-ms epoch (which give rise to the ERP wave) from the modulation of the EEG amplitude across 60 trials.

SVD is a method for decomposing the data matrix **M** *(s x e)*, in this case EEG signals: *s =* 1024 time samples, *e* = 60 trials (given that the sampling rate is 1024 Hz, each 1000-ms epoch has 1024 samples) into *s* wave components *(left singular vectors*, defined as the columns of a matrix **U**(*s x e*)) and *e* habituation components *(right singular vectors*, defined as the columns of a matrix *Y(e x e))*. The left-singular vectors tell us how the EEG amplitude is modulated across the 1000-ms epoch (wave component), and the right-singular vectors describe how the EEG amplitude is modulated across 60 trials (habituation component). Each left-right component pair is multiplied by a scaling factor **σ**, and pairs are rank-ordered according to those factors, where the most important pairs correspond to the largest values of σ, and the least important ones (typically noise) correspond to the lowest σ. Formally, SVD is given by 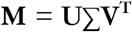, where Σ is a *s x e* the diagonal *(singular value)*, **U** and **V** are the matrices of left and right singular vectors, respectively, and **V^1^** is the matrix transpose of **Y.** The first component pair gives the optimal rank-1 approximation to the original data matrix, in the least square sense. The first two components give the optimal rank-2 approximation, and so on and so forth.

To test the significance of the SVD decomposition, we separated the variance caused by stimulus-evoked activity from other types of variance (noise), and performed the SVD on the noise traces; finally, we tested whether the results of the SVD performed on the noise traces were different from the SVD performed on **M** (which contains a mixture of signal and noise), adapting an approach previously described (Sengupta and Mitra, 1999; Machens et al., 2010).

Specifically, for each subject and condition, we first estimated the residual noise traces η_i_(*s, e*), by taking the average of the differences between the single-subject EEG amplitude *y*_i_ (*s, e*) and group-average EEG amplitude Y_-i_ (*s, e*) (the group average was calculated after excluding subject *i):*

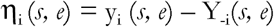

We then performed SVD on the residual noise traces η (*s, e*), for each subject and condition. We averaged the resulting 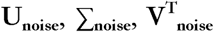 across subjects and divided them by the square root of the number of subjects. We also calculated their standard error of the mean (SEA-1). We tested the significance of the ranks of Σ by comparing whether each diagonal value of Σ was greater than the corresponding value of [Σ_noise_ + 2.33 SE|: this corresponds to a one-tail test at a p-level of 0.01. Lastly, we tested the significance of U and V^1^ by comparing whether their value at each rank was different (either greater or lower) than the corresponding value of [U_noise_ ± 2.58 SEM] and 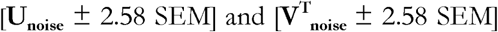: this corresponds to a two-tails test at a p-level of 0.01.

Finally, we modelled the amplitude modulation across trials (habituation components) by fitting the following models to the right-singular vectors at each eigenvalue scale factor (or rank order):

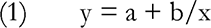

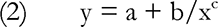

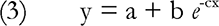

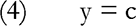

where *y* is the peak amplitude of each given ERP wave, *x* is the trial number (from 1 to 60), *e* is the Euler constant, and *a, b, c* are the parameters to be estimated using a non-linear least squares method. We tested these specific models of ERP decay (#1-3) given the previous evidence that the vertex wave decays sharply at the first stimulus repetition (Fruhstorfer et al., 1970; Greffrath et al., 2007; Mouraux and lannetti, 2009; Valentini et al., 2011; Ronga et al., 2013). Note that model (4) corresponds to the absence of habituation, and fitting this model simply gives c equal to the mean of y. ‘lb compare which model best fitted the data, we calculated the Bayesian Information Criterion (B1C) of each model for each component, ordered by rank, ‘llie B1C allows a fair comparison between models of different complexity because it penalizes models with more parameters (Cover and Thomas, 2006). The lower the B1C, the better the model represents the measured data. For each component rank, we calculated the probability of rejecting the null hypothesis that there was no habituation (i.e., model #4 best represents the data) and accepting the alternative hypothesis that there was significant habituation (i.e., either model #1, 2, or 3 wins), by using a resampling approach with 1000 iterations: at each iteration, we shuffled the order of epochs, fitted models #1-4, and compared the goodness of fit according to B1C.

*4. Code Accessibility*. The code described in “Modelling the within-block decay of the lateralised and vertex waves” was written in Matlab 2016b and is freely available online at [URL redacted for doubleblind review|. The code is available as Extended Data.

## Results

### Response waveforms and topographies

Group-average ERPs elicited by Aβ and Aδ stimuli are shown in Figure. 1, 2 and 3. As expected, the latency of Aδ-ERPs was longer than the latency of Aβ-ERPs, because Aδ fibers are thinly myelinated and thus have slower conduction velocity than large-myelinated Aβ fibers (Mountcastle, 2005).

Figure 1 shows that the amplitude decay of the negative and positive vertex waves (N2 and P2) elicited by the 60 repeated somatosensory stimuli, whereas Figure 2 shows the amplitude modulation of the lateralized somatosensory waves (N1 and P4). To facilitate visual inspection, we enlarged the responses to the first five and last five stimuli (same responses presented both concatenated and super-imposed in figures 1-2). Figure 3 demonstrates that, both in the non-habituated response (trial #1, panels a and c) and in the habituated response (average of trials #6-60, panels b and d), the N2 and P2 waves were greater than baseline. Not only they survived 1-minute of repeated stimulation, but clearly dominated the majority of the ERl’ responses.

In both stimulus modalities, the lateralized somatosensory waves were much smaller than the vertex waves, as expected (Valentini et al., 2012; Hu et al., 2014), and the identification of the Aβ4 peak was ambiguous for the Aβ-ERP elicited by trials 6-60 (figure 3A). Importantly, albeit small in amplitude, both the early N1 and the late P4 lateralized waves elicited in trials 1 and 6-60 were nevertheless consistently greater than baseline, as demonstrated by the point-by-point /-tests reported in Figure 3. The peaks of the N1 waves elicited in trials 1 (panels a and c) and 6-60 (panels b and d) had maximal spatial distribution over the central electrodes in the hemisphere contralateral to hand stimulation (Figure 3), as shown in previous studies (Hu et al., 2014; Mancini et al., 2015).

### Modelling the within-block decay of the lateralised and vertex waves

We took a modelling approach to decompose the modulation of the EEG amplitude across the 1000-ms epoch (which give rise to the ERP wave) from the modulation of the EEG amplitude across 60 trials. This analysis has the benefit of providing an optimal, rank-based approximation to the original data matrix, allowing us to detect habituation effects. Figure. 4 and 5 display the results of the SVD analyses performed at channels Cz (vertex waves) and Cc (lateralised waves) respectively, elicited by non-nociceptive Aβ stimuli (Fig. 4a and 5a) and nociceptive Aδ stimuli (Fig. 4b and 5b). The singular values can be considered as the scaling factors of the left-singular and right-singular vectors. The left-singular vector shows whether and how the EEG amplitude was modulated within the 1000-ms epoch and right-singular vector shows whether and how the EEG amplitude was modulated across 60 trials. The noise distribution for singular, left-singular, and right-singular vectors is shown in red (with 99% confidence intervals). Figure 6 summarises which model best fitted the EEG amplitude modulation across trials, at each rank and according to BIC.

**Figure 4.**
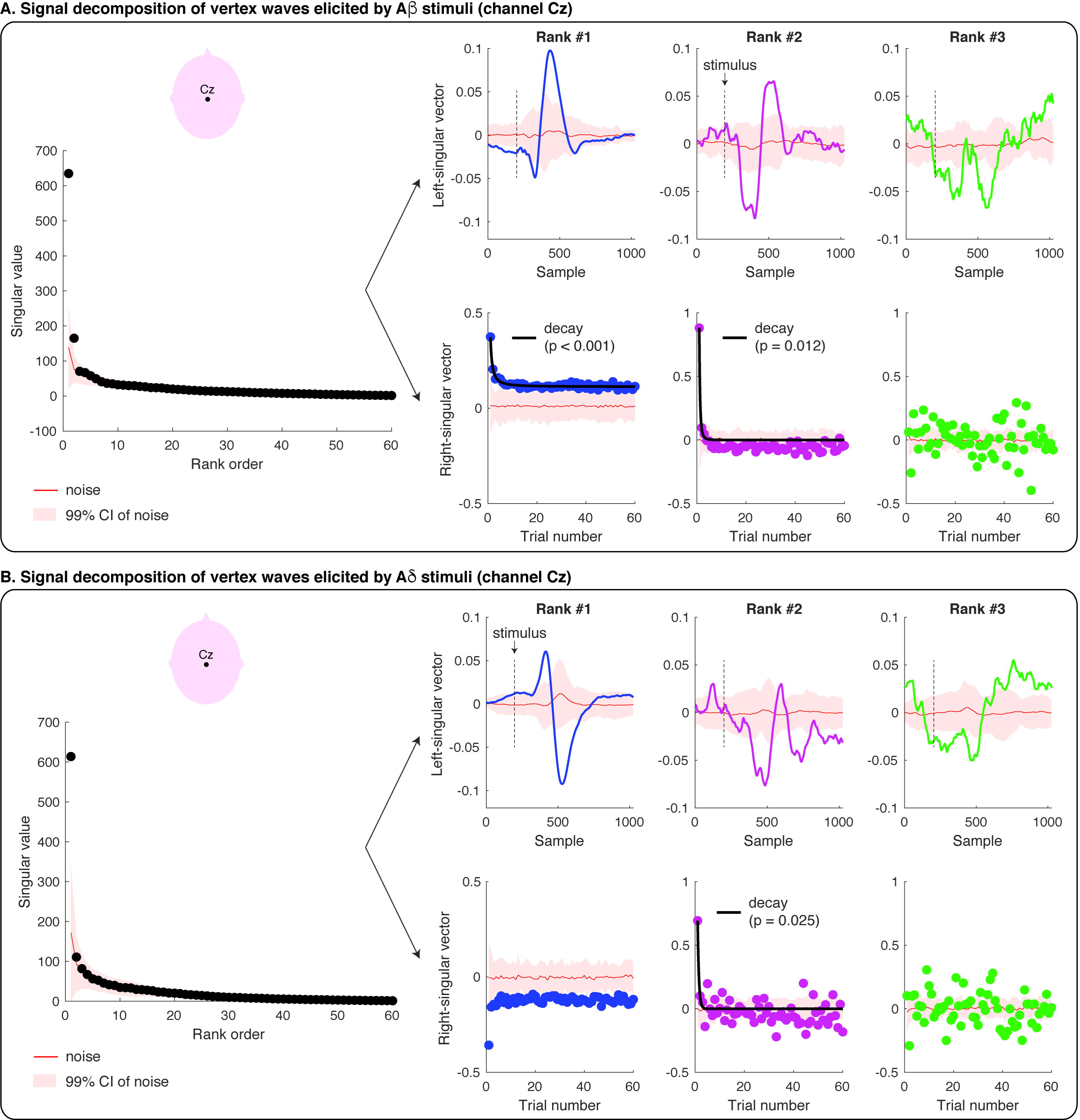
Singular Value Decomposition (SVU) and modelling of the amplitude modulation of the vertex waves (at channel CZ) elicited by repeated Aβ (panel a) and Aδ (panel b) stimuli. Each figure panel displays the singular values at each of the 60 ranks, and the left- and right-singular vectors at the first three ranks. The singular values are the scaling factors of left- and right-singular vectors, and they are ranked according to their importance (from the most important to the least important). The left-singular vector shows the modulation of EEG amplitude across the epoch of 1000 ms (i.e., 1024 samples recorded at 1024 Hz). The stimulus onset is marked with a dashed black line. The right-singular vector shows the modulation of EEG amplitude across the 60 trials. The red line in all plots shows the group-average results of the SVD of the single-subject residual noise traces, with a 99% confidence interval for statistical comparison (p = 0.01). Habituation models were fitted to the right-singular vectors at each rank. If a habituation model wins over a non-habituation model, the fit of the model is displayed with a black line superimposed on the right-singular vector values and the corresponding p-value is reported. In all the instances in which the non-habituation model wins over a habituation model, no fit is displayed.

**Figure 5.**
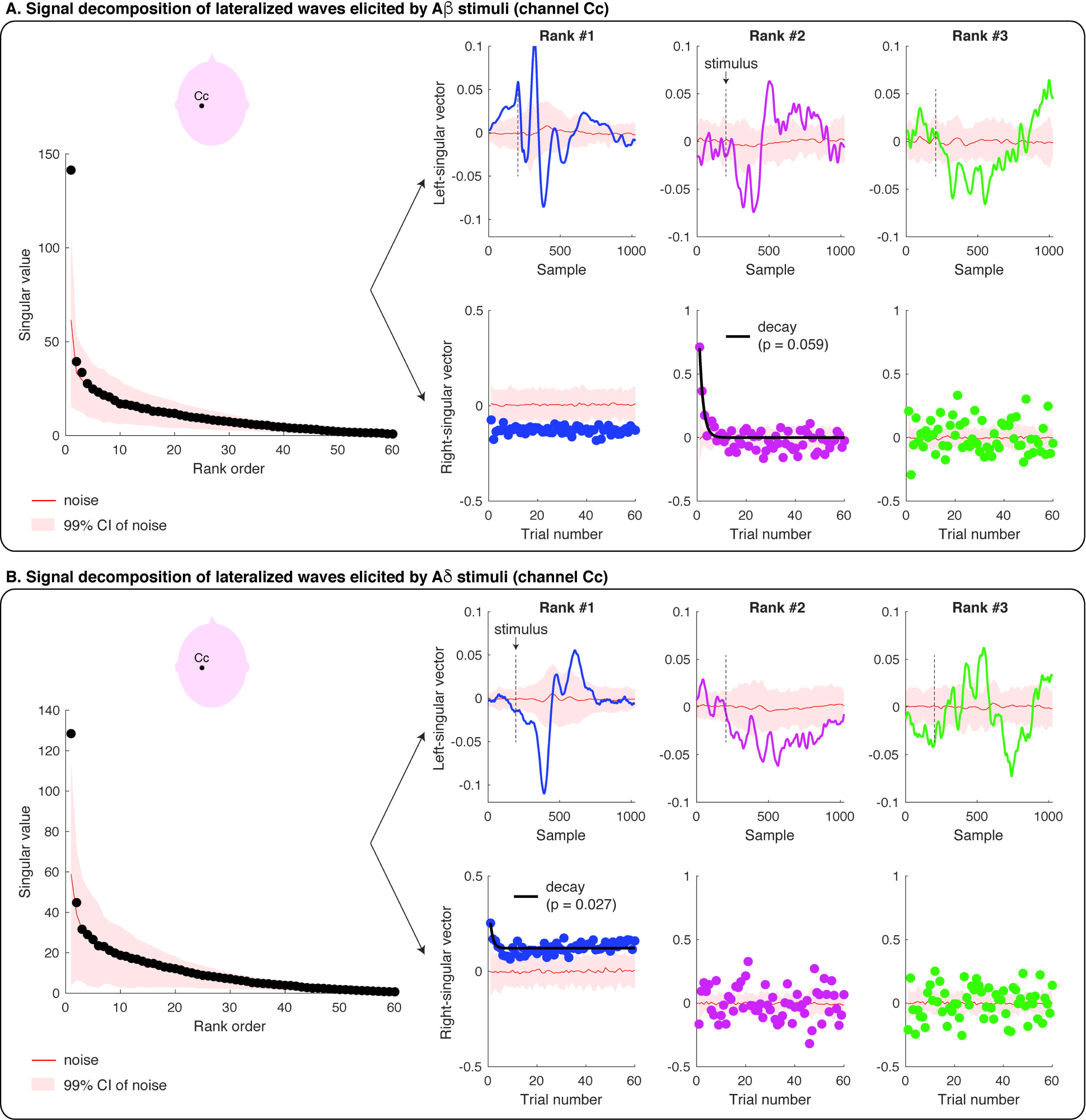
Singular Value Decomposition (SVD) and modelling of the amplitude modulation of the lateralised waves (at channel Cc) elicited by repeated Aβ (panel a) and Aδ (panel b) stimuli. Each figure panel displays the singular values at each of the 60 ranks, and the left- and right-singular vectors at the first three ranks. The singular values are the scaling factors of left- and right-singular vectors, and they are ranked according to their importance (from the most important to the least important). The left-singular vector shows the modulation of EEG amplitude across the epoch of 1000 ms (i.e., 1024 samples recorded at 1024 Hz). The stimulus onset is marked with a dashed black line. The right-singular vector shows the modulation of EEG amplitude across the 60 trials. The red line in all plots shows the group-average results of the SVD of the single-subject residual noise traces, with a 99% confidence interval for statistical comparison (p = 0.01). Habituation models were fitted to the right-singular vectors at each rank. If a habituation model wins over a non-habituation model, the fit of the model is displayed with a black line superimposed on the right-singular vector values and the corresponding p-value is reported. In all the instances in which the non-habituation model wins over a habituation model, no fit is displayed.

The amplitude modulation of the vertex waves elicited by Aβ stimuli was significantly described by the first two ranks (Fig. 4a): the first two singular values were greater than the singular values for the noise distribution (at p-level 0.01). The modulation of the EEG amplitude within the epoch (left-singular vectors) had the characteristic shape of the vertex wave at the first two ranks (Fig. 4a). The latency of the peaks of these waveforms fell clearly within the range of the N2 and P2 peak latencies (Fig. 4a, left-singular vector; cf. Fig. 3a-b): the peaks of the left-singular vector at the first rank had a latency of 125 ms (corresponding to the Aβ-N2 peak) and 225 ms (corresponding to the Aβ - P2 peak); the peaks at the second rank had a latency of 196 ms (corresponding to the late part of the Aβ-N2 wave) and 292 ms (corresponding to the late part of the Aβ-P2 wave). Furthermore, the EEG amplitude elicited by Aβ stimuli decayed significantly across trials at the first two ranks (Fig. 4a, right-singular vector). The winning decay models (*y* = *a+b/ x*) are displayed with a black line superimposed onto the right-singular vectors, and their p-values were <0.001 at rank-1, and 0.012 at rank-2.

The signal decomposition of the vertex waves elicited by nociceptive Aδ stimuli is reported in Fig. 4b. Only the first rank of singular values was greater than noise: at the first rank, the modulation of the EEG amplitude had the characteristic shape and latency of the vertex wave (Fig. 4b, left-singular vector): the peaks of the left-singular vector at the first rank had a latency of 202 ms (corresponding to the peak of the Aδ-N2) and 317 ms (corresponding to the peak of the Aδ-P2; cf. Fig. 3c-d). Although the EEG amplitude clearly decreased from the first to the second trial at the first rank, the fitting of decay models was not significant (Fig. 4b, right-singular vector). Although the second rank of singular values was not significantly different than noise, the modulation of EEG amplitude across time samples was greater than noise at a latency of 270 ms (corresponding to the late part of the N2 wave) and 380 ms (Fig. 4b, left-singular vector): the amplitude of the second-rank component was greater than noise only at the first trial, and its decay was best modelled by the same decay function that described the decay of the vertex wave elicited by Aβ stimuli (*y = a+b/x;* p = 0.025).

The amplitude modulation of the lateralised somatosensory waves elicited by Aβ stimuli (Fig. 5a) and Aδ stimuli (Fig. 5b) was described by the first rank of singular values (p < 0.01). At the first rank, the peak of the left-singular vector fell within the range of the peak amplitude of the N1 wave, both for Aβ stimuli (112 ms) and for Aδ stimuli (181 ms) (Figs. 5a-b; cf. Fig. 3). At the second-rank, the left-singular vector for Aβ stimuli was characterised by two peaks significantly greater than noise: the earliest peak latency fell within the range of the Aβ-N 1 peak latency (112 ms), whereas the second peak had a latency longer than the Aβ-N2 and shorter than the Aβ-P2 peaks (184 ms; cf. Fig. 3a-b). The amplitude of the EEG responses elicited by Aβ stimuli at the first rank was greater than noise (Fig. 5a, right-singular vector), but did not habituate across trials (i.e., the non-habituation model best fitted the right-singular vector). However, at the second rank, the EEG amplitude of the first three trials was greater than noise, and the signal habituation was again in the form of *y = a+b/x* (Fig. 5a, right-singular vector; p = 0.059). Finally, the ERP elicited by Aδ stimuli significantly habituated across trials: indeed, the right-singular vector at the first rank habituated following the same decay functions of the N2 and 1’2 waves elicited by Aβ stimuli and Aδ stimuli (*y = a+b/x*; p = 0.027; see also Fig. 6).

## Discussion

In this study, we characterised the habituation of the different components of the ERPs elicited by 60 identical somatosensory stimuli (activating either Aβ non-nociceptive or Aδ nociceptive primary afferents) delivered at 1 Hz. Although the response amplitude was clearly reduced, the spatiotemporal sequence of the ERP waves was overall preserved in the habituated response (Figure. 3). This was substantiated by point-by-point statistical analysis: both lateralised somatosensory components and supramodal vertex components typically observed in the ERP elicited by sporadic and unpredictable stimuli (Liang et al., 2010; Hu et al., 2014; Mancini et al., 2015) also contributed to the ERP elicited by frequent and predictable stimuli. This result challenges a previous report that 60 repetitions of nociceptive stimuli at 1 Hz fully suppresses lateralised waves (Mouraux et al., 2013) and indicates that lateralised waves are obligatorily elicited by nociceptive-selective stimulation. Furthermore, we used SVD to decompose the modulation of the EEG amplitude across the 1000-ms epoch (which give rise to the ERP wave) from the modulation of the EEG amplitude across 60 trials. We found that the same model described the habituation of the vertex waves and lateralised waves elicited by Aβ and Aδ stimuli (Eigs. 4-6): that was the simplest decay function in the form of *y = a+b/x*, where *y* is the EEG amplitude, .vis the trial number, and *a, b* are the estimated parameters. This indicates that the amplitude of both vertex and lateralised waves decays monotonically, with a largest, transient drop of response magnitude at the first stimulus repetition, followed by much smaller decreases in subsequent repetitions.

### Effect of stimulus repetition on somatosensory lateralised responses

In somatosensory ERPs, the VW is both preceded and followed by other deflections of smaller amplitude. These have a topographical distribution maximal over centro-parietal electrodes in the hemisphere contralateral to hand stimulation. The earliest negative wave is usually referred to as N1 (Valentini et al., 2012) and the latest positive waveform of somatosensory ERPs is referred to as 1’4 (Hu et al., 2014; Mancini et al., 2015). Whereas the 1’4 has only been recently identified and its significance is not yet understood, the N1 has been described repeatedly in a large body of studies (Treede et al., 1988; Spiegel et al., 1996; Garcia-Larrea et al., 2003; Lee et al., 2009; Hu et al., 2014; Mancini et al., 2015), and largely reflect somatosensory neural activities (Lee et al., 2009; Liang et al., 2010).

The neural origin of the N1 wave has been long debated and remains unresolved, but it seems to be at least partially different in the ERPs elicited by non-nociceptive and nociceptive somatosensory stimuli (Garcia-Larrea et al., 2003; Ohara et al., 2004; Frot et al., 2013). A number of studies performing intracerebral recordings have indicated that the Aδ-N1 wave is largely contributed by the operculo-insular cortex (Frot et al., 1999; Peyron et al., 2002; Valeriani et al., 2004), whereas other studies have indicated that both the N1 and 1’4 waves can also be generated in the primary somatosensory cortex, both in human EEG and rodent ECoG recordings (Treede et al., 1988; Valentini et al., 2012; Hu et al., 2014; Jin et al., 2018). For instance, a previous EEG study (Valentini et al., 2012) has demonstrated that the N1 elicited by nociceptive stimulation of the right and left hand have maximum scalp distribution over the central-parietal electrodes *contralateral* to the stimulated side. In contrast, the N1 elicited by nociceptive stimulation of the right and left foot are symmetrically distributed over the central-parietal *midline* electrodes (see also Treede et al., 1988; Jin et al., 2018). These findings are compatible with the somatotopic representation of the body in the primary somatosensory and motor cortex.

A novel result of our study is that these somatosensory N1 and 1’4 responses are detectable not only in the response to the first stimulus, but also in the habituated ERP response, as supported by the statistical assessment of the scalp distribution of the ERl’ response elicited by both the first and the last stimuli of the series (Figure 3). This is important, given that a previous study using trains of intra-epidermal electrical shocks at 1 Hz failed to observe any lateralized response (Mouraux et al., 2013). We note, however, that in this previous study nociceptive afferents were activated using intra-epidermal electrical stimulation, which can cause strong peripheral and perceptual habituation, more significant than for radiant heat stimulation (Mouraux et al., 2010). Thus, in Mouraux et al (2013) peripheral habituation induced by repeated intra-epidermal electrical stimulation in the same skin location may have further reduced the already low signal-to-noise ratio of N1 and P4 waves.

Another novel result of our study is that the lateralised waves habituate across the 60 trials following the same decay functions of the vertex waves (Figure. 4-6). We used SVD not only to decompose the modulation of EEG amplitude within the block and across trials, but also to model the decay of an optimised model of EEG modulation. Indeed, SVD allows separating signals from noise (similarly to Principal Component Analysis) and provides an optimised description of the ERP waves at the most informative ranks. This signal optimization allows characterizing the amplitude modulation of small and noisy ERP components.

**Figure 6.** Winning model of ERP modulation by stimulus repetition. Following singular-value decomposition, three habituation models and a non-habituation model were fitted to the right-singular vectors at each of the 60 ranks and compared according to BIC. The winning models are color-coded (pink: *y — a +b/x*, white: no habituation). Other decay models never win (blue, yellow).

A previous MEG study has reported that neural activity originating from primary somatosensory cortex is more resilient to stimulus repetition (2-Hz pneumatic stimulation of the fingers and face): in other words, it decays to a less extent and more slowly than neural activity in higher-order cortical regions, such as the posterior parietal cortex (Venkatesan et al., 2014). We used slower stimulus frequencies than these studies, so we cannot exclude that different time-scales of habituation may emerge at faster stimulus repetitions.

Finally, our design was not suited to investigate the habituation of the earliest sensor) components of Ap-ERPs, which typically require averaging responses elicited by hundreds of stimuli. However, we note that the N20 wave of Aβ-ERPs, which originates in area 3b, is very resilient to stimulus repetition (Garcia Larrea et al., 1992) and is not modulated by selective spatial attention (Garcia-Larrea et al., 1991). In contrast, the later N1 waves of Aβ- and Aδ-ERPs can be modulated by spatial attention (Legrain et al., 2002).

### Effect of stimulus repetition on vertex ERI responses

The negative-positive vertex wave (VW) is the largest component of the EEG response elicited by sudden sensor) stimuli. Converging evidence indicates that stimuli of virtually all sensor) modalities can elicit a VW, provided that they are salient enough (Liang et al., 2010). It is therefore not surprising that the VW elicited by auditor) stimuli repeated at 1-Hz decays following a function similar to the one observed here for somatosensory stimuli (Fruhstorfer et al., 1970). Even when considering experimental observations that did not formally model the response habituation, the maximum decrease in VW amplitude consistently occurs at the first stimulus repetition, for auditor) (Ritter et al., 1968; Fruhstorfer et al., 1970), somatosensory (Larsson, 1956; Fruhstorfer, 1971; Iannetti et al., 2008; Wang et al., 2010; Valentini et al., 2011; Ronga et al., 2013) and visual stimuli (Courchesne et al., 1975; Wastell and Kleinman, 1980). The similarity of the decay of the VW elicited by Aβ and Aδ stimuli (Figure. 1, 3, 4) further supports the multimodal nature of the neural generators of these signals (Mouraux and Iannetti, 2009). The mechanisms underlying such sharp reduction of response amplitude at the first stimulus repetition are likely to be similar across sensor) systems.

Before discussing the contribution of the present results in elucidating the functional significance of the VW, it is important to highlight the empirical evidence that the observed response habituation is not due to neural refractoriness of afferent neurons or to fatigue of primary receptors. A previous study recorded ERPs elicited by pairs of nociceptive stimuli delivered at short intervals, which could be either identical or variable across the block (Wang et al., 2010). Only when the inter-stimulus interval was *constant* across the block, the VWs elicited by the second stimulus were reduced in amplitude. The peak amplitude of the VWs elicited by the second stimulus was instead as large as the VWs elicited by the first stimulus when the inter-stimulus interval was *variable*, indicating that neither neural refractoriness nor fatigue can easily explain the sharp response decay to stimulus repetition.

Furthermore, if the sharp response habituation at the first stimulus repetition was determined by fatigue of primary sensory receptors, we would have observed different decay profiles for stimuli delivered in varying vs constant spatial locations. Indeed, the VW elicited by contact heat stimuli at long and variable intervals (8-10 seconds) decays much faster if the second stimulus is delivered at the same spatial location of the first (Greffrath et al., 2007). Instead, we observed remarkably similar patterns of ERP decay for both Aδ laser stimuli delivered at different spatial locations and A β electrical stimuli delivered in the same skin region. Additionally, electrical stimuli activate directly the axons in the nerve trunk, bypassing the receptor, further ruling out receptor fatigue as explanation for the Aβ-ERP habituation. Receptor fatigue might still contribute to the slow decrease in ERP magnitude observed across dozens of stimulus repetitions of laser stimuli (Greffrath et al., 2007), but certainly not to the dramatic reduction of ERP amplitude we observed after one single stimulus repetition.

The physiological significance of the VW remains to be properly understood. However, there is evidence that this large electrocortical response reflects neural activities related to the detection of salient environmental events (jasper and Sharpless, 1956; Mouraux and Iannetti, 2009) and execution of defensive movements (Moayedi et al., 2015; Novembre et al., 2018). The detection of salient events relies on a hierarchical set of rules that consider both their probability of occurrence and their defining basic features (Legrain et al., 2002; Wang et al., 2010; Valentini et al., 2011; Ronga et al., 2013; Moayedi et al., 2016). The present results are informative with respect to this functional framework. Indeed, stimulus repetition did not abolish the VW elicited by either Aβ or Aδ stimuli, although it reduced its amplitude already after the first stimulus repetition. Therefore, even when stimulus saliency is reduced by contextual factors, there is a residual activity of the VW generators, only minimally reduced after the first few stimulus repetitions (Figure. 1, 3b, 3d). These findings point towards the existence of an obligator) VW activity triggered by any sudden and detectable change in the environment, even when contextual modulations minimize its behavioural relevance.

Extensive evidence from cell physiology indicates that neural habituation to repeated stimuli arises from alterations of synaptic excitability. Even the simple gill-withdrawal reflex in Aplysia dramatically habituates at the first stimulus repetition (Byrne et al., 1978), due to a decreased drive from the sensor) neurons onto follower motor neurons (Castellucci et al., 1970; Carew and Kandel, 1973). The temporal profile of this short-term habituation follows a fast decay function (Carew and Kandel, 1973), strikingly similar to that observed in this and other studies on the habituation of electrocortical responses in humans (Fruhstorfer et al., 1970; Greffrath et al., 2007). These synaptic changes have been interpreted as a hallmark of learning, and are central to the ability of the nervous system to adapt to environmental events (Carew and Kandel, 1973). Interpreting the decay of neural responses as functionally relevant for learning is not in contradiction with attentional interpretations: stimuli that are learned and recognized are likely to require less attentional resources than novel stimuli, and stimuli that need to be learned are typically more salient.

### Conclusion

In conclusion, our results provide a functional characterization of the decay of the different ERP components when identical somatosensory stimuli are repeated at 1Hz. Nociceptive and non-nociceptive stimuli elicit ERPs obligator) contributed by both lateralised and vertex components, even when stimulus repetition minimizes stimulus relevance. This challenges the view that lateralised waves are not obligatorily elicited by nociceptive stimuli. Furthermore, the lateralised and vertex waves habituate to stimulus repetition following similar decay functions, which most possibly cannot be explained in terms of fatigue or adaptation of skin receptors.

## Acknowledgments

FM and GDI were supported by a Wellcome Trust Strategic Award (COLL JLARAXR). GDI is additionally supported by a ERC Consolidator Grant (PAINSTRAT) and by the Medical Research Council. AM is supported by an ERC Starting Grant (PROBING-PAIN). The authors declare no competing financial interests.

